# Semantic Influences on Object Detection: Drift Diffusion Modeling Provides Insights Regarding Mechanism

**DOI:** 10.1101/2024.06.24.600369

**Authors:** Jingming Xue, Robert C. Wilson, Mary A. Peterson

## Abstract

Research shows that semantics, activated by words, impacts object detection. Skocypec & Peterson (2022) indexed object detection via correct reports of where figures lie in bipartite displays depicting familiar objects on one side of a border. They reported 2 studies with intermixed Valid and Invalid labels shown before test displays and a third, control, study. Valid labels denoted display objects. Invalid labels denoted unrelated objects in a different or the same superordinate-level category in studies 1 & 2, respectively. We used drift diffusion modeling (DDM) to elucidate the mechanisms of their results. DDM revealed that, following Valid labels, drift rate toward the correct decision increased, i.e., SNR increased. Following Invalid labels, SNR was lower only for upright displays in study 2. Thresholds were higher in studies 1 & 2 than in control. That more evidence must be accumulated from displays that follow labels implies that familiar object detection entails semantic activation. Threshold was even higher following Invalid labels in study 2, suggesting that more evidence from the display is needed to resolve within-category conflicts. These results support the view that semantic networks are engaged in object detection.

## Introduction

A long-standing question regarding visual perception is whether knowledge such as the meaning of words (i.e., semantics) can influence how we perceive objects (Abdel Rahman & Sommer, 2008; Costello et al., 2009; Fodor, 1984; Koffka, 1935; Meteyard et al., 2007; Noorman et al., 2018; Pinto et al., 2015; Rosenthal, 2006; Sander, 2006; Stein & Peelen, 2015). For example, does receiving information about a stimulus beforehand make it easier to *recognize* it later? Previous work suggests that, for object **recognition**, the answer is yes (Boutonnet & Lupyan, 2015; Carr et al., 1982; Dils & Boroditsky, 2010; Rabovsky et al., 2012; Samaha et al., 2018). Recent research has suggested that object **detection** – considered a more basic perceptual process than recognition – is also influenced by semantics (e.g., Lupyan & Ward, 2013; Maier & Abdel Rahman, 2018; Weller et al., 2018). These detection findings might suggest that high-level semantics influence perception far down the perceptual hierarchy. However, controversy remains over whether those previous studies assessed object detection *per se* or the detection of features of the denoted objects (Skocypec & Peterson, 2022). Moreover, if semantics does influence object detection, questions must be answered regarding how it does so. Whereas experimental results can support some hypotheses, modeling can further elucidate mechanism.

In the present article, we use computational modeling to reveal the mechanisms whereby semantics influenced object detection in a recent experiment by Skocypec and Peterson (2022). We begin by discussing the “Label-Feedback” hypothesis, a leading hypothesis regarding how semantics might influence object detection that was supported by previous experiments in which stimuli were repeated multiple times. In the next section, we discuss an alternative theoretical interpretation supported by Skocypec and Peterson’s (2022) results. We discuss Skocypec and Peterson’s manipulations and results in some detail before introducing drift diffusion modeling, the modeling method used in this article.

### A Leading Hypothesis: The Label-Feedback Hypothesis

The “Label-Feedback” hypothesis regarding how semantics might influence object detection suggests that presenting a valid label for an object shortly before the object appears (e.g., the word “umbrella” before an image of an umbrella) activates representations of that object all the way down the visual hierarchy to low-level features. This pre-activation improves object detection by enabling faster progression through the visual hierarchy from lower-level features to higher-level object representations (Lupyan, 2012; Lupyan & Clark, 2015). On this hypothesis, an invalid label would interfere with object detection by activating low-level features of a different object; mismatching input would necessitate revising those predictions which would slow progression through the visual hierarchy. In studies that provided support for the Label-Feedback hypothesis, a basic-level label (i.e., a label commonly paired with an object) was paired with a particular exemplar many times (e.g., Lupyan & Ward, 2013; Pinto et al., 2015; Stein & Peelen, 2015). However, with so much repetition, participants may have learned to respond accurately following a basic-level label when they detected low-level features rather than the object configuration which entails representations of the relationship among features (Gayet et al., 2014; Skocypec & Peterson, 2022). Thus, the use of repetition may produce a misunderstanding of how basic-level labels typically affect object detection.

### An alternative Semantic Network hypothesis

Skocypec and Peterson (2022) proposed an alternative account of how basic-level word labels affect object detection: Without repetition, valid basic-level labels activate a distributed semantic network that represents words, objects, object properties, object affordances, commonly performed actions, etc. (e.g., see Clarke & Tyler, 2014; Martin, et al, 2018). Object representations in this network take the form of neural populations that are tuned by previous encounters with the objects (Ashbridge et al., 2000; Gibson & Peterson, 1994; Perrett et al, 1998; Peterson, 2019); hence, more units are tuned to the typical, upright, configuration of the represented object than to atypical orientations. Therefore, activity in a neural population accumulates more quickly when an upright rather than an upside down, “inverted”, object is encountered. If decisions about the presence of an object depend on accumulated evidence in the neural population selective for that object configuration per se rather than its features, then detection decisions would be faster and/or more accurate for upright than inverted objects (e.g., Peterson & Gibson, 1994; Stokes et al., 2009). In contrast, detection decisions based on features should not vary with orientation because features alone do not activate configurations. Therefore, one way to determine what type of representations mediate semantic effects on object detection is to investigate whether effects of labels depend on the orientation of the object denoted by the word. If low-level feature representations are responsible, the effects of word labels should be orientation independent. In contrast, if via a semantic network, the words activate neural populations representing the objects they denote, then the effects of word labels should be larger for upright than inverted objects.

### Skocypec and Peterson’s (2022) study

Skocypec and Peterson (2022) assessed object detection via figure assignment reports regarding stimuli like those in Figure 1. In these stimuli, a portion of a well-known (i.e., “familiar”) object with a typical upright is depicted on one side of a central border that divides the display into two equal-area halves. The term “familiar” in this article refers to previous experience outside the laboratory with objects in the same basic-level category, not to repetition within the experiment. The objects depicted in the bipartite stimuli were new instances of well-known objects. They were not repeated in the experiments. In previous research Peterson and colleagues had found that figures (i.e., objects) are more likely to be detected on the side of the border where a familiar configuration is sketched when the display is upright than when it is inverted. For example, participants are more likely to perceive the object on the left in Figure 1A where a portion of a woman is depicted in a familiar upright configuration rather than in Figure 1B when the woman is depicted in an unfamiliar inverted orientation; the features are the same, but the configuration is different. Likewise, the figure/object is more likely to be detected on the right in Figure 1C than Figure 1D (Peterson, 2019). This orientation dependency implicates input to object detection decisions from neural populations representing the objects in these displays.

**Figure 1.**
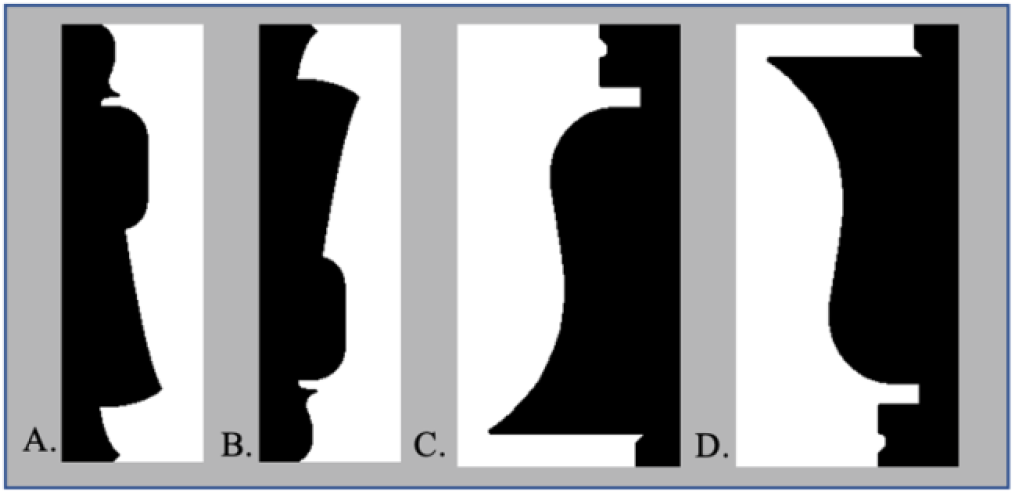
Examples of bipartite stimuli. A portion of a well-known object was sketched on one side of the central border of stimuli in which two equal-area regions, one black and one white lay on opposite sides of that border. One, “critical”, region depicted a well-known object in an upright or an inverted orientation and was equally often on the left and right, in black and white. A. The critical region depicting a portion of a woman in an upright orientation is on the left in black. B. An inverted version of A. C. The critical region depicts a portion of a bell in an upright orientation in black on the right side. D. An inverted version of C. Black/white color of the critical regions was balanced in the experiments.

In their experiments, Skocypec and Peterson (2022) investigated the effect of word labels on object detection in these bipartite displays. Participants in experimental labels-present groups were presented with either a valid or an invalid label before being shown a brief, masked, exposure of a bipartite display depicting a portion of a familiar object in either an upright or an inverted orientation. Valid labels denoted the object at a basic level (e.g., valid labels for the objects in Figure 1 were “woman” and “bell”).

Skocypec and Peterson leveraged the orientation dependence of familiar configuration effects to examine whether effects of valid word labels shown before bipartite displays are mediated by activation of low-level features or by higher-level neural populations representing configured objects. If low-level features drive the effect, label effects should be orientation invariant; if higher-level neural populations are key, label effects should be larger for upright than inverted objects. To be noted, individual participants viewed each bipartite display (upright or inverted) and each label once only to assess semantic influences unaffected by repetition.

Valid labels preceded the bipartite displays on half the trials; invalid labels denoting unrelated objects preceded the displays on the other half of the trials (label type was randomly intermixed). To ensure the words are unrelated to the objects in the target display, we used the database SUBTL Word Frequency Database (Brysbaert & New, 2009). The invalid labels used in study 1 and study 2 were different. In study 1, the invalid labels denoted an object from a **different** superordinate-level category, where the superordinate-level categories were natural versus artificial objects. For instance, the invalid labels for woman, a natural object, and bell, an artificial object, were, respectively, “money” (artificial) and “fish” (natural). In study 2, the invalid labels denoted an object from the **same** superordinate-level category as the object sketched in the upcoming display, albeit unrelated (e.g., “shark” (natural) and “book” (artificial) for woman and bell, respectively). To determine whether valid labels improved object detection, invalid labels impaired object detection, or whether both effects occurred, Skocypec and Peterson also compared performance in these labels-present studies to performance in a control study in which the same displays were not preceded by words (labels-absent control).

Overall, Skocypec and Peterson’s (2022) results supported the hypothesis that effects of basic-level word labels shown once only are best explained by activation in semantic networks rather than by predictions regarding the low-level features of objects: In both studies, valid labels improved detection accuracy and response times (RTs) over control and effects were larger when the familiar objects in the bipartite stimuli were depicted in an upright rather than an inverted orientation (Figure 2). Furthermore, in study 1, different superordinate-level category invalid labels did not affect detection accuracy or response times compared to results obtained in the control labels-absent study. These results were also contrary to predictions from a Label-Feedback account where interference is expected from invalid predictions. In study 2, however, following same superordinate-level category invalid labels detection RTs were substantially and significantly increased compared to control for both upright and inverted objects. This result was unexpected on the Label-Feedback hypothesis because fewer prediction revisions should be necessary when invalid labels denote objects in the same rather than a different superordinate-level category. Skocypec and Peterson attributed these results to a conflict in the overlapping semantic networks activated by the invalid label and the object in the test display, given that objects in the same superordinate-level category are highly likely to share properties, features, potential actions, etc. (cf., Martin et al., 2018). Skocypec and Peterson hypothesized that this conflict had to be resolved before detection occurred.

**Figure 2.**
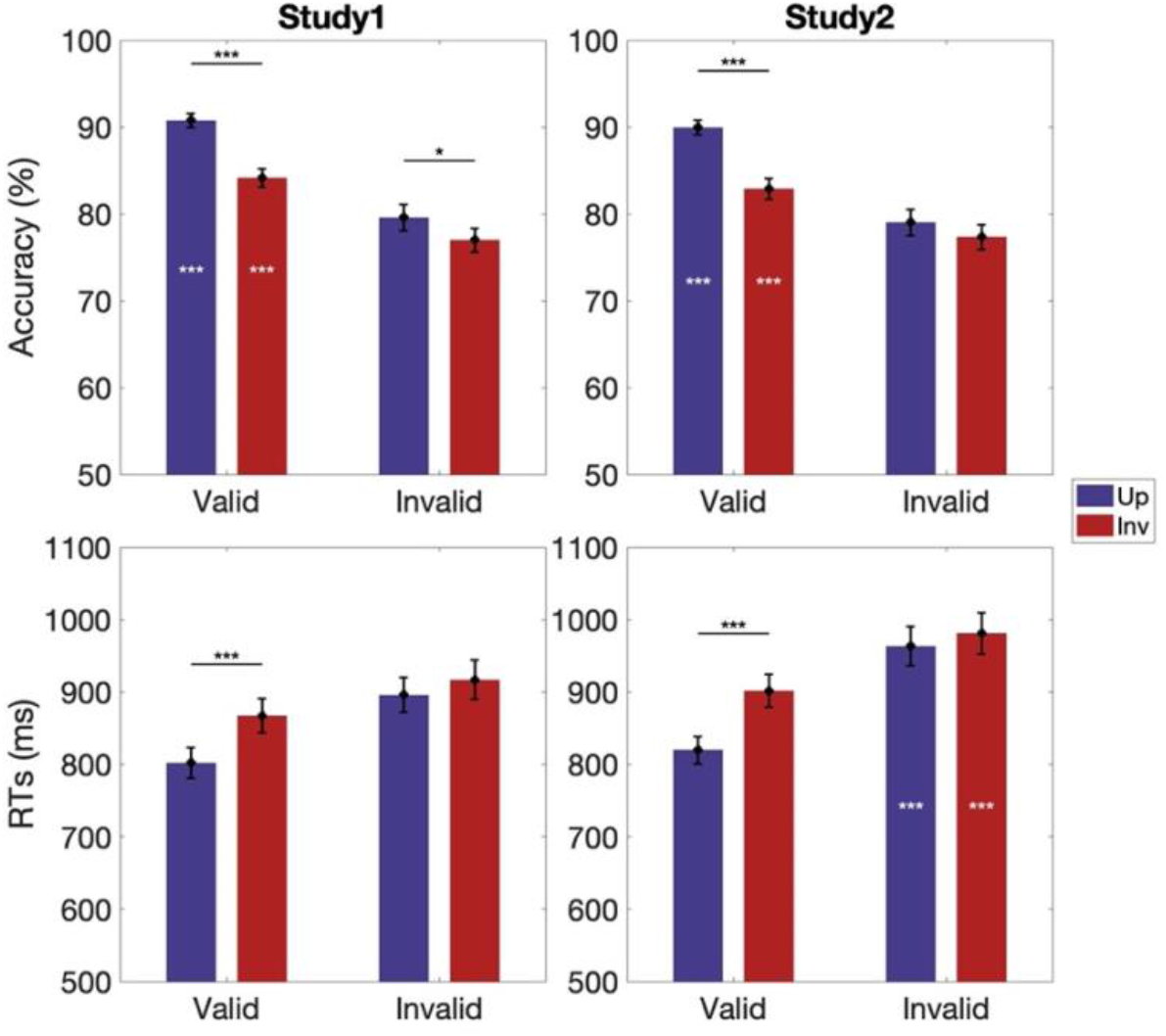
Skocypec and Peterson’s (2022) behavioral results. (First column) study 1; (Second column) study 2. (First row) Accuracy; (Second row) response times. Blue bars: Performance with upright displays; Red bars: Performance with inverted displays. White asterisks indicate main effects of labels-present vs. labels-absent groups. Horizontal lines and black asterisks indicate orientation-dependent differences. Error bars represent pooled standard errors. *** indicates p < 0.001, * indicates p < 0.05.

Skocypec and Peterson interpreted their result as evidence that object detection is not simply affected by semantic activation; it *entails* semantic activation. This was a new proposal. Here we use computational modeling to elucidate the mechanisms of Skocypec and Peterson’s results.

### The Present Research

We used a drift diffusion model (Bogacz et al., 2006; Ratcliff, 1978; Figure 3) to reveal the underlying cognitive computations involved in detecting the objects in the bipartite displays in Skocypec and Peterson’s (2022) experiments. By jointly modeling object detection accuracy and RT on participants’ individual trials (rather than mean RTs per participant per condition like Skocypec and Peterson), this model can tell us whether the object detection decision is controlled by a change in *drift rate* (roughly processing speed), a change in evidence *threshold*, or a change in starting point (here, a left/right side bias). Drift diffusion models have proven valuable for studying neural mechanisms across a wide range of species and contexts (e.g., Bogacz et al., 2010; Feng et al., 2021; Lositsky et al., 2015; Pedersen et al., 2017; Ratcliff et al., 2004; Tavares et al., 2017; Wilson & Collins, 2019), with their parameters known to associate with neuronal activity (e.g., Gold & Shadlen, 2007a).

**Figure 3.**
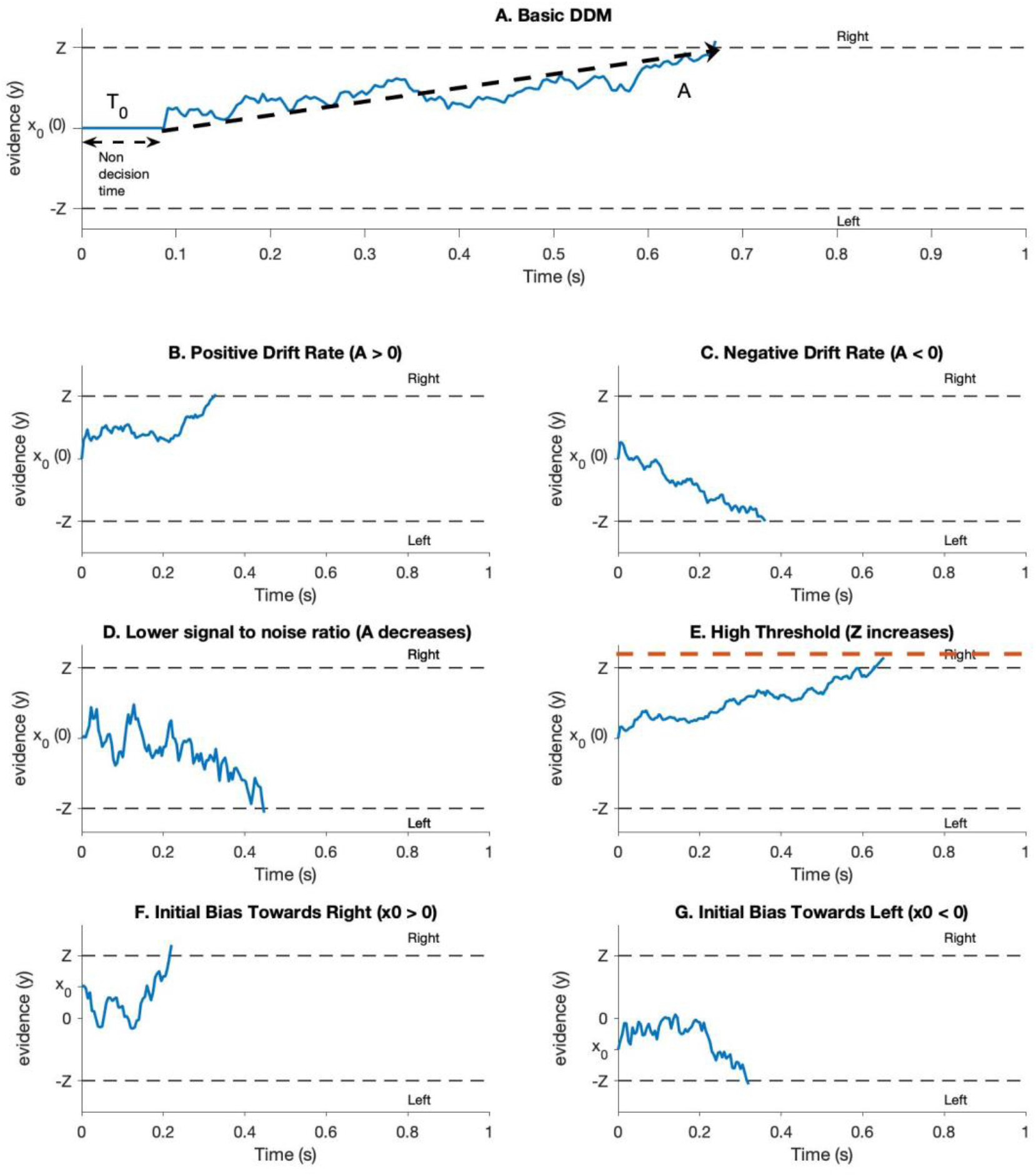
Different cases of drift diffusion model. (A) A basic drift diffusion model over time. The x-axis represents time in seconds, and the y-axis represents evidence accumulation, which could be a cognitive or a perceptual process leading towards a decision. The dashed lines at the top and bottom represent decision thresholds for two choices, labeled as "Right" and "Left", denoting by 𝑍 and −𝑍 separately. Evidence accumulates over time, starting from 𝑥0 (in this case, 0) and fluctuating until it reaches one of the thresholds, indicating a decision has been made. The double headed arrow denotes the non-decision time, 𝑇0 . In A, the evidence crosses the upper threshold, suggesting a decision made towards the "Right side" option. (B) A decision process with a positive drift rate, A, on evidence accumulation (A > 0). A positive drift rate indicates a tendency for the evidence to accumulate towards the "Right side” choice over time. (C) A negative drift rate (A < 0) indicates a tendency to accumulate evidence towards the “Left side" choice. (D) A lower signal-to-noise ratio where there is more variability and noise in the evidence accumulation process than in C, resulting in a longer path to a decision threshold. (E) A higher threshold condition where more evidence must be accumulated before decision. (F) Evidence accumulation begins with a bias towards the "Right" choice, starting above zero, indicating a predisposition towards that option. (G) The opposite initial bias towards the "Left" choice, with evidence accumulation starting below zero.

Our drift diffusion model (DDM) assumes that, when presented with a bipartite test display, participants make a left/right response by accumulating “evidence” over time in favor of one response or the other. This evidence is assumed to contain both signal – whether the object lies to the left or right of the central border – as well as noise caused e.g., by ambiguous cues and random neuronal responding.

Thus, the accumulated evidence, 𝑦, evolves according to a drifting random walk with a noise parameter, 𝑐, controlling the randomness of the walk and a drift parameter, 𝐴, controlling the drift (see Figure 3A).

In Figure 3B, a positive drift rate denotes evidence that the object is on the right and the evidence tends to drift upwards over time. Conversely, a negative drift rate corresponds to the case in which the object is on the left; here, evidence tends to drift down (Figure 3C). The relative magnitude of the drift rate to the noise (i.e., |𝐴|/𝑐) denotes the “signal-to-noise ratio,” with a higher ratio implying stronger evidence and faster processing and a lower ratio implying weaker evidence and slower processing (Figure 3D). An initial condition, 𝑥_0_, controls the initial bias for one option or the other (e.g., a bias for responding right/left, Figures 3F and G).

The model makes a decision when the evidence crosses one of two “decision thresholds” at 𝑦 = ±𝑍.

In Figures 3B and 3C, the model chooses “right side” when it crosses the positive threshold and “left side” when it crosses the negative threshold. The time at which the model crosses the threshold is known as the Decision Time (DT), the time taken by the decision process. This Decision Time is further related to the Response Time (RT) through the addition of a non-decision time 𝑇_0_, which captures sensory, motor, and other non-decision related delays between the onset of the sensory stimulus and the response. Thus, by modeling which threshold is crossed the DDM can model the accuracy, and by modeling when the thresholds are crossed the DDM can model the response times. By fitting the model to human choices and response times, we can therefore fit the four free parameters of the model – drift rate, 𝐴 ; the initial condition, 𝑥_0_; the threshold, 𝑧; and non-decision time, 𝑇_0_ – and investigate how they change in different label conditions. The model includes an additional noise parameter that is fixed at a value of 1. This parameter is integrated with the drift rate to determine the signal-to-noise ratio. For further details, please refer to the methods section.

### Mapping the drift Diffusion model to Skocypec and Peterson’s (2022) results

By identifying the parameters of the label effects in a DDM, we aim to test Skocypec and Peterson’s (2022) Semantic Network interpretation of their results. First, we can test their hypothesis that the orientation effects following valid labels occur because the valid label operates through a semantic network to pre-activate the neural population representing the familiar object it denotes. On this hypothesis, the drift diffusion model should return a higher drift rate for the upright than the inverted condition when valid word labels precede the displays compared to the control condition. Second, we can assess the mechanisms of the different invalid label effects observed in study1 and study 2. Are the differences due to changes in drift rate or changes in threshold or both? We will investigate whether there is study- or condition-dependent starting point differences to investigate whether participants have an a priori bias to press the right or left button.

## Method

We applied our DDM to the response times for both accurate and inaccurate responses recorded by Skocypec and Peterson (2022). In what follows, we briefly describe their method in more detail.

### Participants

Data were analyzed from 321 undergraduate students (18–36 years old; M = 19.21, SD = 1.85; 236 F and 85 M) from the University of Arizona (UA) who participated for course credit or payment. The total included participants who participated in one of 12 experiments: an original and a replication for each of two exposure durations (90 ms and 100 ms) in either the control no-labels study; study 1 (a labels- present study with different superordinate-level category invalid labels intermixed with valid labels); or study 2 (a labels-present study with same superordinate-level category invalid labels intermixed with valid labels)). Skocypec and Peterson (2022) found no substantive differences between the two exposure conditions; hence the results are combined here. All participants provided informed consent and had normal or corrected-to-normal vision. (For more information about participants see Peterson and Skocypec (2022).

### Test displays/Stimuli

The stimuli were bipartite displays, in which a central border divided a vertically-elongated rectangle into two equal-area regions (samples shown in Figure 1), one black and one white. The central border sketched a portion of a well-known object (*a familiar configuration*) on one side; this was the "critical side". The critical side was equally often on the left/right and colored black/white. The stimuli were shown on a medium gray backdrop. All three studies used the same 72 displays, including 36 upright and 36 inverted displays. In upright displays, the critical side sketched a familiar object its typical upright orientation. The inverted displays were obtained by rotating the upright display 180° and mirroring them on the vertical axis. Half of the familiar configurations depicted portions of natural objects, and the other half depicted portions of artificial objects. The side complementary to the critical side depicted a novel shape. (For normative data regarding these stimuli, see Flowers et al., (2020).) Every participant saw each display once only. Stimuli were balanced across conditions in 16 programs, each seen by eight participants per study. More information about the displays can be found in Peterson and Skocypec (2022) and at https://osf.io/j9kz2/.

### Procedure

Participants were provided with a consent form approved by the Human Subjects Protection Program at the University of Arizona. After they signed it, an experimenter tested their visual acuity using a Snellen eye chart. Next, participants were introduced to the object detection task; they were told that their task was to report whether they detected a shaped entity -- a figure – on the left or right side of the central border by pressing a left (right) button on a response box. They made their response with the index and middle fingers of their dominant hand. They were instructed to report their first impression of where they detected a figure. This was our assay of object detection. Participants were also informed that there were no correct or incorrect responses; that we were interested in what they saw. Responses made up to 4000 ms after test display onset were recorded.

The test procedure is shown in Figure 4. Each trial started with a central fixation cross. Participants were instructed to press a foot pedal when they were ready for the trial to begin. In studies 1 and 2 (the labels-present studies), after the foot pedal was pressed, a label was shown for 250 ms followed by a blank screen. Next, a bipartite display was shown briefly; different groups viewed the displays for 90 or 100 ms. (Skocypec and Peterson (2022) found that exposure duration did not differentially affect accuracy in the labels-present studies versus the control study. Hence, the present analysis uses data from the two exposure durations.) The bipartite display was followed immediately by a mask (200 ms) and then by a blank screen that was present until response or 4000 ms had elapsed. Participants’ RT was recorded from the onset of the bipartite display. Half of the labels in studies 1 and 2 were valid in that they were basic-level category labels for the object sketched on the critical side of the central border in the upcoming display. The other half were invalid in that they denoted a different object. Both studies used the same valid labels. The invalid labels were different in the two studies: In study 1 the invalid labels denoted an unrelated object from a **different** superordinate-level category from the object in the display; in study 2, the invalid labels denoted an unrelated object from the **same** superordinate-level category as the object in the display. The procedure was the same in the control study except that no word labels were presented before the bipartite test displays. (See Figure 4A.)

**Figure 4.**
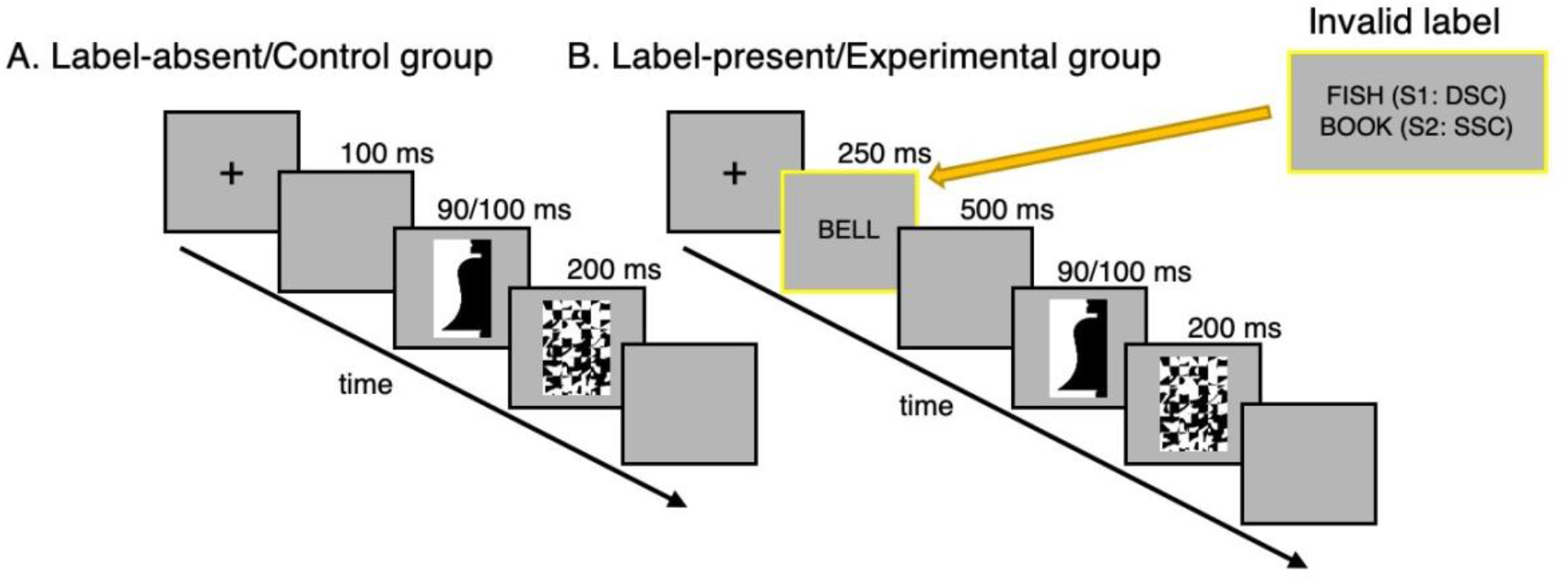
Trial structures for labels-present and labels-absent studies. (A) Trial structure for labels absent experiments (control studies). Each trial began with a fixation point shown on a medium gray background (which was used throughout), followed by a 100 ms blank screen. Then, the stimulus was presented for 90 ms or 100 ms to different groups, followed by a mask for 200 ms and then by a blank screen. (B) Trial structures for labels present experiments, the critical difference from control is that, after the fixation cross, a word label appeared in the center of the screen for 250 ms. The label was either valid, denoting the object in the upcoming bipartite display at a basic level, or invalid, denoting a different unrelated object. Invalid labels in study 1 denoted an object in a different superordinate-level category (i.e., natural vs. artificial), denoted by "DSC" in the figure; invalid labels in study 2 denoted an object in the same superordinate-level category, denoted by "SSC" in the figure. The inset shows sample invalid labels for the object depicted in the bipartite display (i.e., bell) in study 1 (S1) and study 2 (S2). For a complete listing of the invalid labels see the Appendix in Skocypec and Peterson (2022).

### Drift Diffusion Model

#### The basic drift diffusion model

The basic drift diffusion model (DDM) models decision making between two options as an evidence accumulation process (Figure 3A). In this view, the model integrates noisy information about the decision over time to form a decision variable 𝑦 that captures the relative evidence for option 1 (e.g., right) vs option 2 (e.g., left). 𝑦 > 0 denotes that the model has more evidence in favor of option 1, 𝑦 < 0 denotes that the model has more evidence in favor of option 2. During the decision process, the evidence starts at some initial value, 𝑥_0_, that captures the initial bias in favor of one option of the other, if any, and evolves over time according to the following stochastic differential equation:

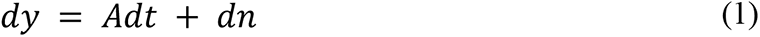

where 𝑑𝑦 denotes the instantaneous change in evidence, 𝑑𝑡 denotes a small unit of time, 𝐴 denotes the drift rate (the average signal in favor of one option or the other) and 𝑑𝑛 denotes Gaussian random noise of mean 0 and variance 𝑐^2^𝑑𝑡. Simulating Equation 1 leads to the evidence taking a drifting random walk over time, where the variance of the walk is governed by the noise parameter 𝑐 and the drift by the drift rate 𝐴. It is therefore typical in the DDM literature to set the noise parameter 𝑐 = 1 and interpret the drift rate 𝐴 as a signal-to-noise ratio (i.e., Setting the noise parameter *c* to 1, the ratio *A/c* simplifies to *A*. With c standardized to 1 across different studies, the drift rate *A* can now be directly interpreted as the signal-to-noise ratio, which makes the model more interpretable and standardize comparisons across studies, this is a normalization step that simplifies the model without changing its fundamental characteristics).

The DDM makes a decision when it crosses one of two thresholds, choosing option 1 if it crosses the top threshold at +𝑧 and option 2 if it crosses the bottom threshold at −𝑧. The time at which it crosses the threshold corresponds to the decision time (DT), which is related to the response time (RT) via

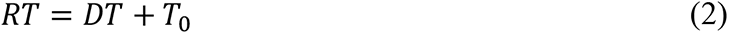

where 𝑇_0_ is a parameter of the model corresponding to the amount of time not related to the decision – e.g. sensory processing time, motor preparation time.

Overall, the DDM has five free parameters: the initial condition, 𝑥_0_, the drift rate, 𝐴, the noise, 𝑐, the threshold, 𝑍, and the non-decision time 𝑇_0_. Ideally, we would fit all five parameters to the behavioral data. Unfortunately, however, because some of these parameters only appear as ratios in expressions for accuracy and response time, only four parameters can be independently estimated from data (Bogacz et al., 2010). Thus, as stated above, we set the noise parameter 𝑐 = 1 and interpret the drift rate 𝐴 as a signal-to-noise ratio. To apply this model to our tasks, we next had to make assumptions about how these parameters might change in different task conditions.

#### Application of the drift diffusion model to the control experiment

First, we applied the DDM to the simpler control experiment. In this experiment, familiar objects can appear on the left or right side of the border (side, 𝑠 = +1 for right, -1 for left) and in an upright or inverted orientation (orientation, 𝑜 = +1 for upright, 0 for inverted). We assumed that side and orientation could affect the drift rate in the model, specifying an equation for the drift rate as

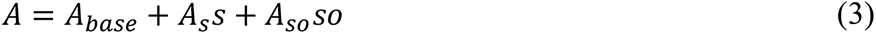

Where 𝐴_𝑏𝑎𝑠𝑒_ describes the baseline drift rate corresponding to a drift bias to left or right, 𝐴_𝑠_ denotes the main effect of side and 𝐴_𝑠𝑜_ denotes how the effect of side is modulated by orientation.

Similarly, we assumed that side and orientation could also impact the starting point, writing the initial condition as:

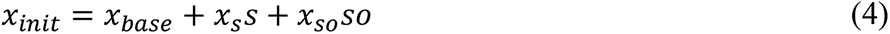

Where 𝑥_𝑏𝑎𝑠𝑒_ describes the initial bias to left or right, 𝑥_𝑠_ denotes the main effect of side and 𝑥_𝑠𝑜_ denotes how the effect of side is modulated by orientation.

Finally, we assumed that the threshold 𝑍 and non-decision time 𝑇_0_, were constant regardless of the side and orientation of the stimulus. This assumption, that the threshold and non-decision time are independent of the stimulus, is common in drift diffusion models of behavior and reflects the idea that threshold is set before the stimulus is presented (Bogacz et al., 2006; Wilson & Collins, 2019).

Thus, for the control task, our model has 8 free parameters that we fit to the responses and response times of the participants: 𝐴_𝑏𝑎𝑠𝑒_, 𝐴_𝑠_, 𝐴_𝑠𝑜_, 𝑥_𝑏𝑎𝑠𝑒_, 𝑥_𝑠_, 𝑥_𝑠𝑜_, 𝑍, and 𝑇_0_.

#### Application of drift diffusion model to main task

To extend our model to capture the effects of word labels, we also included the effect of the word label type (𝑙 = +1 for valid, 0 for invalid) on the drift rate, starting point, and threshold. Thus, for the main task, the equation for drift rate becomes:

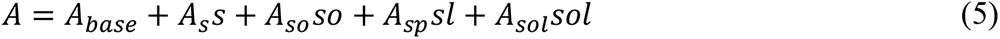

Where 𝐴_𝑠𝑝_ denotes the change in drift rate when a valid word label is present and 𝐴_𝑠𝑜𝑙_ denotes the change in drift rate when a valid word label is present and the stimulus is upright (As mentioned before, *l* = 0 for invalid).

Similarly for the starting point we have

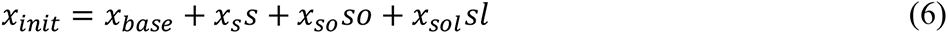

Where 𝑥_𝑠𝑜𝑙_ denotes the change in starting point when a valid word label is present and the stimulus is upright.

Because the label appears before the stimulus, we assume that it can affect the threshold, writing

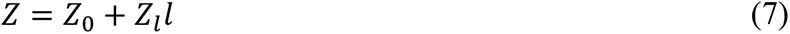

Where 𝑍_0_ denotes the baseline threshold and 𝑍_𝑙_ the change in threshold for a valid label (Again, *l* = 0 for invalid).

Finally, we assumed that the non-decision time was constant regardless of labels.

Thus, our model for the task has 13 free parameters, 𝐴_𝑏𝑎𝑠𝑒_, 𝐴_𝑠_, 𝐴_𝑠𝑜_, 𝐴_𝑠𝑙_, 𝐴_𝑠𝑜𝑙_, 𝑥_𝑏𝑎𝑠𝑒_, 𝑥_𝑠_, 𝑥_𝑠𝑜_, 𝑥_𝑠𝑜𝑙_, 𝑍_0_, 𝑍_𝑙_ and 𝑇_0_.

### Model fitting

We fit the DDM to behavior from the Skocypec and Peterson (2022) studies using a maximum likelihood approach. Each participants’ data were fit separately to the model to account for variability in cognitive processing across individuals. This method allowed us to adjust the parameters of the DDM (𝐴, threshold 𝑍, starting point 𝑥_0_, and non-decision time 𝑇_0_) for each participant to maximize the likelihood of observing the actual response times and accuracy rates recorded in the experiment. The fitting process was implemented in Matlab using the fmincon function. Data and code to replicate these analyses are available at https://osf.io/g3mxb/.

With so many free parameters in the model an obvious concern is that they will not be identifiable from data. To test whether this was the case, we performed a parameter recovery analysis (Wilson & Collins, 2019). The results, shown in the Appendix, showed an excellent correspondence between the fitted and ground truth value for each parameter suggesting that all parameters are identifiable in our task. Therefore, we further conducted parameter comparisons across different conditions to investigate the effect of semantic labels.

## Results and Discussion

### Basic behavior

As can be seen in Figure 5, after fitting, the drift diffusion model captures all the main effects in both the control and experimental studies. In the control labels-absent studies, the model captures the more accurate responses in the upright condition (i.e., the orientation effect) without any orientation-dependent RT differences. In both labels-present studies, the model captures the orientation effect and faster and more accurate responses following valid labels. Thus, at least qualitatively, the model captures all the main effects in the data. We next examine the evidence regarding condition-dependent drift rate, evidence threshold, and starting point to reveal the mechanisms of Skocypec and Peterson’s (2022) results.

**Figure 5.**
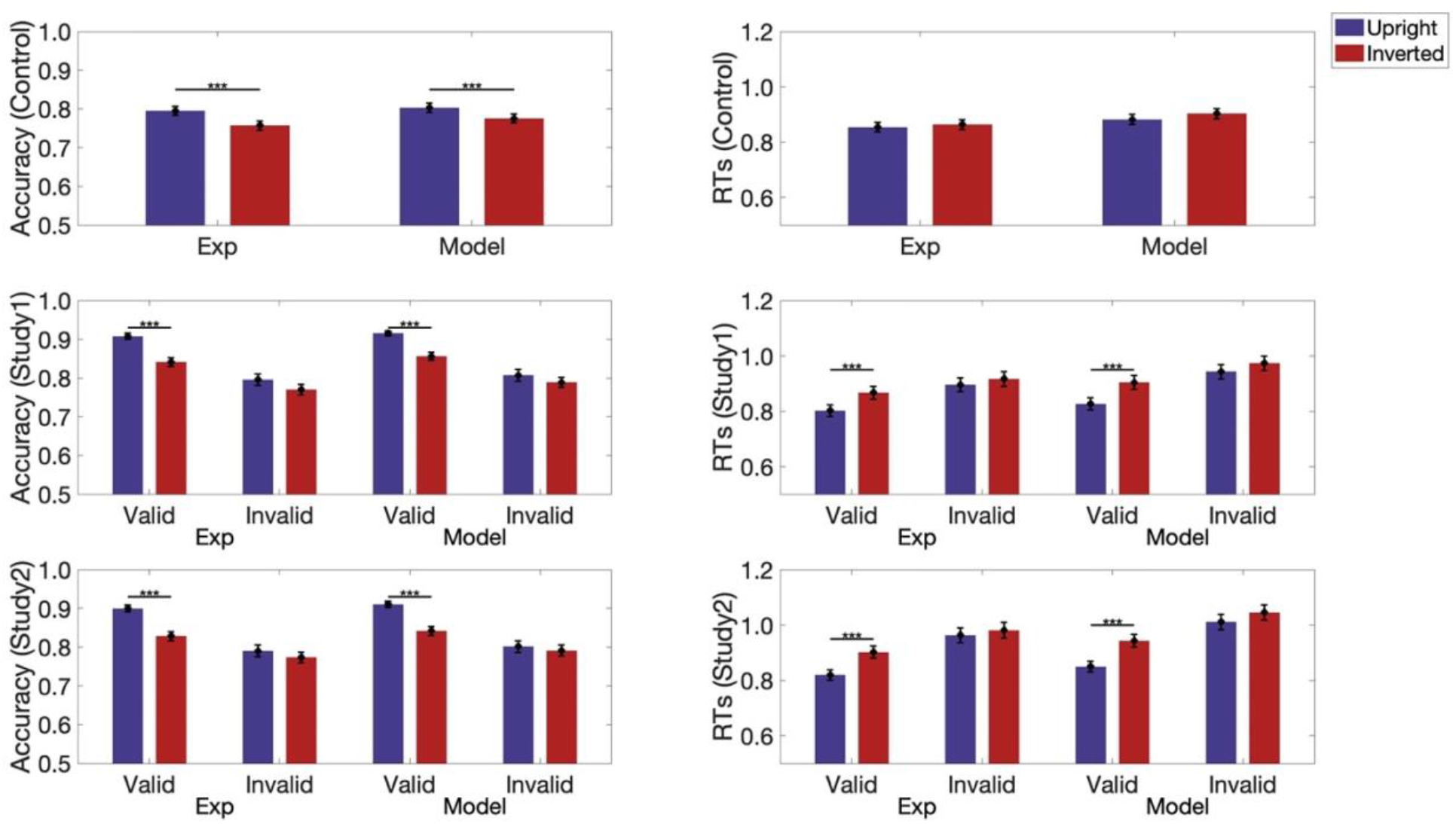
Accuracy (column 1) and Response times in seconds (column 2) for both experimental data and data simulated from the model for both control (row 1) and experimental studies 1 and 2 (rows 2 & 3). Blue bars represent the upright condition. Red bars represent the Inverted condition. Error bars represent pooled standard errors. *** indicates *p* < 0.001.

### Drift<u> rates</u>

Drift rates obtained in the labels-present studies (studies 1 and 2) were first compared to drift rates obtained in the control labels-absent study to determine whether valid labels increase drift rate, invalid labels reduce drift rate, or both effects occur. Specifically, for each of the studies, a 2 (study: 1 [or 2] vs. control) x 2 (orientation: upright vs. inverted) mixed ANOVA was conducted for drift rates following valid and invalid labels separately. To preview: our drift diffusion model results show that valid labels increase drift rate; they also show that invalid same category labels can reduce the fastest drift rates, but invalid different category labels did not affect drift rates. As discussed below, both effects are best explained by the Semantic Network hypothesis.

#### Valid labels versus No Labels

In both study 1 and study 2, drift rates were significantly higher when valid labels preceded the displays compared to when no labels preceded the displays in the control study and the increase was larger for the upright than inverted displays (see Figure 6, top row). For both studies these effects were evident in main effects of orientation and study and an interaction between orientation and study: Drift rates were higher in the upright condition than the inverted condition: for study 1, *F*(1, 216) = 45.84, *p* < 0.001, *η2* = 0.18 and for study 2, *F*(1, 214) = 41.41, *p* < 0.001, *η2* = 0.16.

**Figure 6.**
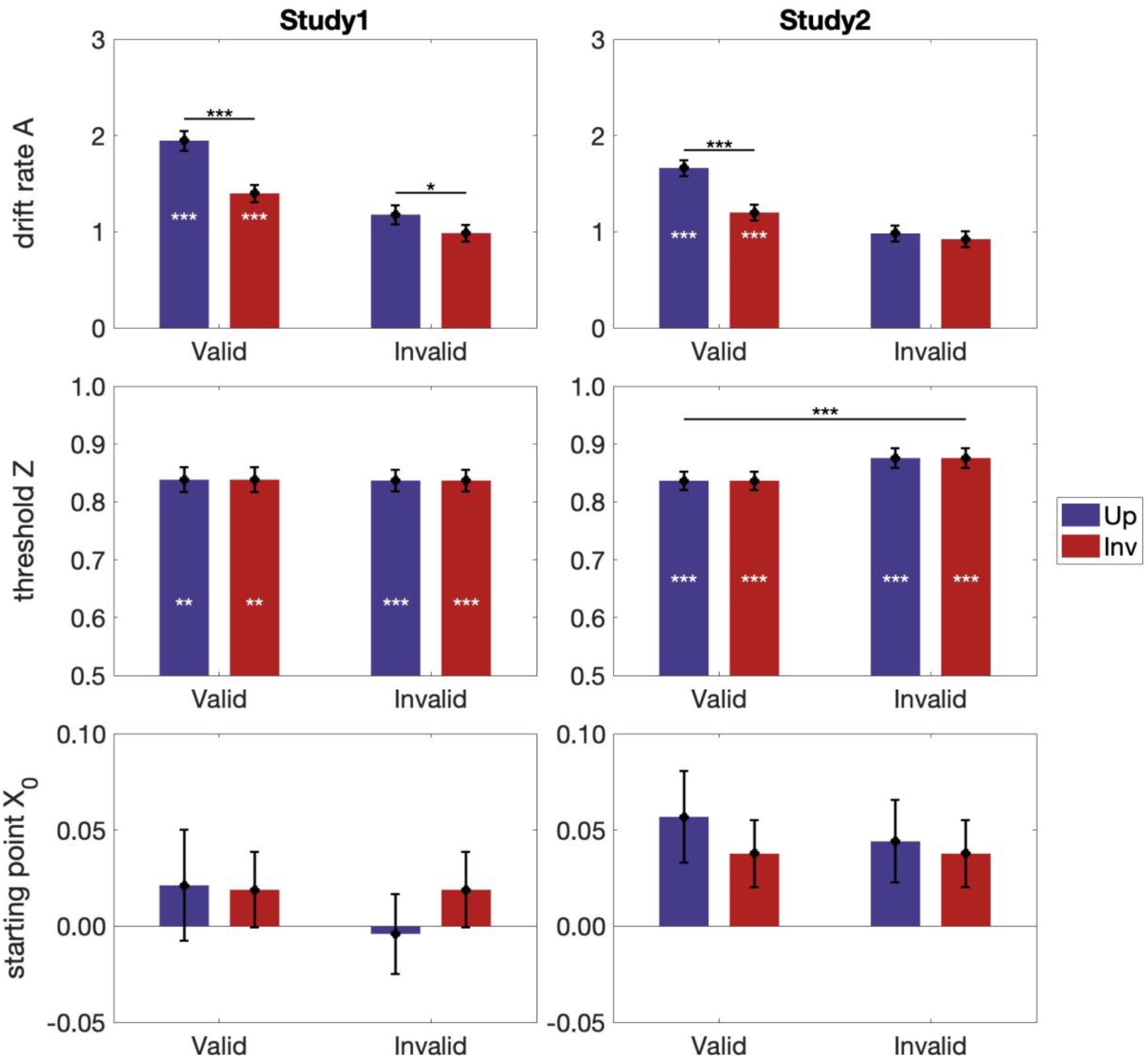
Model parameters in labels-present studies. (First row) drift rate (*A*) (Second row) Threshold (*Z*). (Third row) Starting point (𝑥_0_). Higher values indicate a greater drift rate/threshold/starting point. For starting point, positive values indicate starting points toward the right side, negative values indicate starting points toward the left side. (Left column) study 1; (Right Column) study 2. Blue: upright displays; Red: inverted displays. White asterisks indicate statistically significant experimental vs. control differences. Error bars represent pooled standard errors. *** indicates *p* < 0.001, ** indicates *p* < 0.01, and * indicates *p* < 0.05

Drift rates were higher in the labels-present studies than in the control study: for study 1, *F*(1, 216) = 42.96, *p* < 0.001, *η2* = 0.17 and for study 2, *F*(1, 214) = 21.16, *p* < 0.001, *η2* = 0.09. For both studies, the interaction between study and orientation reached significance, for study 1 *F*(1, 216) = 7.27, *p* = 0.008, *η2* = 0.03 and for study 2, *F*(1, 214) = 4.37, *p* = 0.038, *η2* = 0.02. Simple effects analyses indicated that the increase in drift rate following a valid label was larger in the upright than in the inverted condition: Control drift rates were 1.086 for upright displays and 0.85 for inverted displays. Study 1 mean difference upright = 0.860 (SD for the mean difference = 0.134, 95%CI [0.597, 1.124], *p* < 0.001); mean difference inverted = 0.548 (SD for the mean difference = 0.109, 95%CI [0.333, 0.763], *p* < 0.001). The same was true for study 2: mean difference upright = 0.576 (SD for the mean difference = 0.123, 95%CI [0.334, 0.818], *p* < 0.001); mean difference inverted = 0.349 (SD for the mean difference = 0.105, 95%CI [0.142, 0.556], *p* = 0.001). The evidence that drift rate is higher following valid labels in the labels- present studies than in the control labels-absent study and that the increase is greater for upright than inverted displays support the Semantic Network hypothesis rather than the Label-feedback hypothesis where orientation effects are not predicted.

#### Comparison of drift rates following valid labels in study 1 and study 2

To examine whether drift rates on valid label trials were affected by the context shaped by the invalid labels shown on other trials in the labels-present studies, we conducted a 2 (study: 1 vs. 2) x 2 (orientation: upright vs. inverted) mixed ANOVA on valid drift rates. Drift rates following valid labels were higher in study 1 (*M* = 1.672) than in study 2 (*M* = 1.431), *F*(1, 206) = 5.08, *p* = 0.025, *η2* = 0.02 (see Figure 6, first row). This small, but statistically significant effect indicates that the rate of evidence accumulation following valid labels is affected by the context established by the type of invalid labels present on other trials within a study. In study 2, the semantic network of the target object overlaps with the semantic network of the object denoted by invalid labels because they are both in the same superordinate-level category. This overlap, present on 50% of the trials, may introduce enough noise in the evidence accumulation process to cause the reduction in drift rate observed on valid trials.

#### Invalid labels versus No Labels

Mixed ANOVAs comparing drift rates following invalid labels to those observed in the control labels-absent study were conducted separately for study 1 and 2. The factors were study (1 [or 2] vs. control) and orientation (upright vs. inverted). For both studies, this comparison revealed main effects of orientation: drift rates were higher when displays were upright rather than inverted (study 1: *F*(1, 216) = 18.67, *p* < 0.001, *η2* = 0.08; study 2: *F*(1, 214) = 11.41, *p* = 0.001, *η2* = 0.05). (See Figure 6, top row.) For study 1, orientation effects were observed in both the control labels- absent study and in the labels-present study; no interaction between study and orientation was observed, *F*(1, 216) = 0.20, *p* = 0.653. For study 2, however, an interaction was observed between orientation and study (*F*(1, 214) = 4.02, *p* = 0.046, *η2* = 0.02): An effect of orientation was present in the control study but not in study 2: control: mean upright – inverted difference = 0.236 (SD = 0.061, 95%CI [0.117, 0.355], *p* < 0.001); study 2: mean upright – inverted difference = 0.060 (SD = 0.063, 95%CI [-0.065, 0.185], *p* = 0.344). Thus, it seems that the fastest drift rates -- those typically found for upright objects – are reduced by the same-category invalid labels in study 2. This result is likely due to noisy evidence accumulation following invalid labels in study 2 as discussed in the previous section. This pattern of results is not consistent with the Label-Feedback hypothesis, where the rate of evidence accumulation would be expected to be lower when invalid labels denote objects with different rather than similar features (i.e., in study 1 rather than study 2). An interpretation in terms of noise due to overlapping semantic activation by the label and the object present in the test display can also account for threshold effects as we discuss next.

### Threshold

The model was designed such that evidence thresholds would be the same for upright and inverted stimuli. This is because participants are unaware of the objects’ orientation until after object detection (i.e., Participants were not required to recognize the objects, only to detect them. Indeed, in the brief masked exposures they may not have recognized the objects or noted the correspondence between upright and inverted stimuli). For the control study, the evidence threshold was 0.760.

We next examined the evidence thresholds obtained in studies 1 and 2 following valid and invalid labels separately. The mean threshold following valid labels was 0.838 in study 1 and 0.837 in study 2. These thresholds were statistically higher than the thresholds in the control study; for study 1, *t*(104) = 3.16, *p* = 0.002; for study 2, *t*(102) = 3.62, *p* < 0.001. (Figure 6, middle panel.) That more evidence is required following valid labels in the labels-present studies than in the control labels-absent study indicates that before a detection decision is made, semantic activation initiated by the object in the display must be distinguished from semantic activation initiated by labels, at least when valid and invalid labels are intermixed as they are in studies 1 and 2. In turn, this result suggests that semantic activation initiated by the familiar configuration in the test displays contributes to object detection responses when labels are absent in the control study; otherwise, there would not be an increase in threshold following valid labels in studies 1 and 2. In other words, these results are consistent with Skocypec and Peterson’s (2022) proposal that the detection of well-known objects like those depicted in the bipartite displays is not just *affected by* semantic activation; it *entails* semantic activation.

To examine whether the different types of invalid labels in study 1 versus 2 differentially affected thresholds, we conducted a 2 (study: 1 vs. 2) x 2 (label type: valid vs. invalid) repeated measures ANOVA on thresholds. There were no main effects of label type, *F*(1, 206) = 3.53, *p* = 0.062, or study, *F*(1, 206) = 0.60, *p* = 0.440 for study. An interaction between study and label type was observed, *F*(1, 206) = 4.14, *p* = 0.043, *η2* = 0.02. Simple effects analyses showed a higher threshold following invalid than valid labels in study 2: threshold on invalid label trials = 0.876; mean difference = 0.039 (SD for the mean difference = 0.014, 95%CI [0.011, 0.067], *p* = 0.006) but not in study 1: threshold on invalid label trials = 0.837; mean difference = 0.002 (SD for the mean difference = 0.014, *p* = 0.912). The interaction with study suggests that more evidence must be accumulated from the display before a detection response can be made when the semantic network of the object denoted by an invalid label immediately preceding the test display overlaps with the semantic network of the object in the display. The hypothesis that object detection entails semantic activation is supported by the evidence that response thresholds were higher following invalid labels than valid labels in study 2, where the invalid labels denoted objects in the same- superordinate level category as the object on the critical side of the border, but not in study1, where the invalid labels denoted objects in a different-superordinate level category, .

### Starting Point

We compared the starting point in the control study to the starting point obtained in studies 1 and 2 following valid and invalid labels separately using Mixed ANOVAs. The factors were study (1 [or 2] vs. control) and orientation (upright vs. inverted). These analyses failed to reveal any differences in starting point in either study. No significant main effects or interactions were found in these analyses (Figure 6, third row). For study 1, *p*s > .129; for study 2, *p*s > 0.189 (see appendix for details). These null results are not unexpected; participants were not expected to prefer one of the two sides before evidence accumulation began because the object was depicted equally often on the left and right sides of the displays.

## Discussion

Our goal in the present study was to elucidate the mechanisms underlying the results of Skocypec and Peterson’s (2022) investigation of whether valid and invalid labels shown before test displays affect object detection per se and not feature detection. To do so, we applied a drift diffusion model (DDM) to individuals’ response times on all trials regardless of whether their responses were accurate or inaccurate. Recall that Skocypec and Peterson had analyzed individuals’ mean responses on accurate trials only. The DDM derived three potential mechanisms for how words affect object detection: (1) By changing the drift rate, i.e., the rate at which evidence for the detection response accumulates (i.e., processing speed); (2) By changing the evidence threshold, i.e., the the detection boundary representing the amount of evidence that must be accumulated from the display before a detection response is made; (3) By changing the starting point for a decision, i.e., whether one of two potential responses is favored at the outset. Analyzing the full set of results, we found that, compared to the control studies without labels, labels influenced object detection through altering both drift rate and threshold. No changes in startinig point were observed.

*Drift Rate*. Compared to a control labels-absent study, valid labels shown before test displays increased the rate of evidence accumulation for the object in the display; the increase was larger for upright than for inverted objects. Invalid labels had very little inlfuence on behavior. This is exactly the pattern predicted if words activate a semantic network that includes the neural population representing the denoted object. We propose that the prior activation of the test object’s neural population reduces the number of reentrant cycles needed to localize the object to the right or left of the central border (see discussion of Figure 7 below).

**Figure 7.**
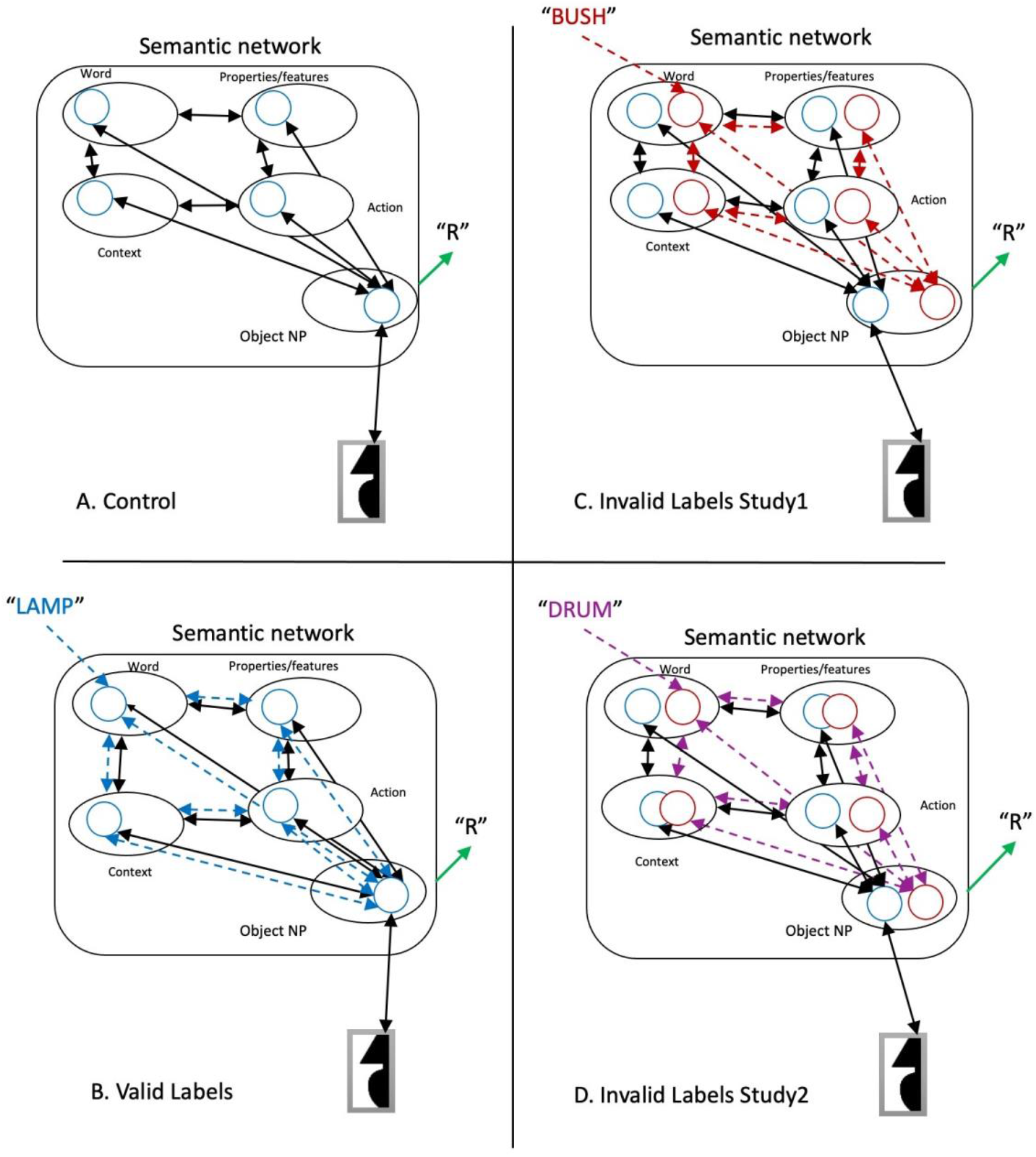
Potential mechanisms operative in Skocypec and Peterson’s (2022) object detection task. Solid black lines with double-headed arrow endings indicate recurrent activity initiated by the object in the bipartite test display, both within the semantic network and between the semantic network and a lower-level representation of the test display (shown below the semantic network). Dashed lines with double-headed arrow endings indicate recurrent activation in the semantic network initiated by labels shown before the test displays, blue for valid labels, red 7 purple for invalid labels. Blue circles in the semantic network indicate semantic representations of the object in the test display in the various portions of the semantic network (e.g., context, object, etc.). Red & purple circles indicate semantic representations of objects denoted by invalid labels. Object NP = neural population representing an object. A) control labels-absent study. B) Valid labels in labels-present studies. (C-D) Invalid labels in labels-present studies; different superordinate-level invalid label as in study 1. D) same superordinate-level invalid label as in study 2.

Valid labels that denoted the object in the test display at a basic level (i.e., bell, woman, etc.) were present on half of the trials in the labels-present studies. On the other half of the trails, invalid labels denoted unrelated objects. In study 1, invalid labels denoted objects in a **differen**t superordinate-level category than the object in the test display. In study 2, invalid labels denoted objects in the **same** superordinate-level category as the object in the test display. The drift rate following valid labels was affected by the context provided by the type of invalid labels shown on the other half of the trials: it was lower in study 2 in the context of invalid labels denoting an object in the same, rather than a different, superordinate-level category as the test object as in study 1. We propose that activating another object in the same superordinate-level category as the target object introduces noise into the evidence accumulation process because the semantic networks of the two objects overlap. This noise, present on 50% of the trials, lowered the signal to noise ratio and resulted in an overall reduction in drift rate. This reduction in drift rate may also account for a second effect of invalid labels: Following same, but not different, superordinate-level invalid labels, drift rates for upright objects were not higher than for inverted objects; this might reflect a reduction in the highest drift rates.

### Threshold

Evidence thresholds were elevated in studies 1 and 2 relative to the control study for both valid and invalid label trials; an interpretation in terms of semantic network noise also accounts for these threshold changes: These results indicate that when labels activated semantics before the appearance of the bipartite test displays, more evidence had to be accumulated from the display to reach the detection threshold than when labels were absent. In both study 1 and study 2, thresholds were elevated compared to the control study even when the labels were valid. This finding implies that object detection responses in the control study were supported at least in part by semantic activation initiated by the object in the display. In the labels-present experiments, however, given that the labels shown before the displays predicted the object in the test display on only 50% of the trials, the mere presence of semantic activation alone could not increase certainty that a particular familiar object was present. Sufficient evidence had to be accumulated from the display to distinguish the semantics of the test object from the semantics of the object denoted by a label shown before the display. In study 1 equivalent threshold increases were observed following valid and invalid labels that denoted objects in a different superordinate-level category from the object in the display. Thus, no additional response uncertainty was induced by the prior activation of different superordinate-level category objects. In study 2, however, thresholds were elevated more following same superordinate-level category invalid labels than valid labels. This finding indicates that more evidence was needed to distinguish the semantics of the object present in the test display from that of an object in the same superordinate category where there is a greater degree of overlap in their semantic networks. It takes time to accumulate this additional evidence, and detection response times are longer. Thus, threshold indices obtained through modeling support Skocypec and Peterson’s (2022) interpretation that object detection entails semantic activation.

Figure 7 illustrates the semantic network implicated by the DDM in label effects on the detection of meaningful objects. The illustration uses a sample test display that depicts a portion of a table lamp on the right side of the central border of a bipartite test display; hence, the accurate response is “Right.” Note that object detection responses, indicated by a green arrow and “R”, are dominated by recurrent activity between the object representation in the semantic network and a lower-level representation of the object in the display, albeit influenced by activity in the semantic network. Panel A illustrates how object detection decisions are made in the control labels-absent study: The critical region in the bipartite display initiates activity in the neural population (NP) representing the object in the display (here, a lamp); see the blue circle in the Object NP portion of the semantic network. The NP includes more representations of the object in its typical, or upright, orientation than in an inverted, upside down, orientation and therefore accumulates evidence faster for upright than inverted objects(Perrett et al., 1998). As evidence accumulates in the Object NP, activity spreads to other components of the semantic network associated with that object (e.g., words denoting that object, object properties and features, contexts in which that object is likely to occur, actions related to the object, etc.); see the blue circles in other parts of the semantic network. Reentrant activity within the semantic network enhances activity throughout the network (shown by double-headed black arrows in the semantic network). Reentrant processes from the Object NP to lower retinotopic areas are necessary to localize the object on one side of the border (see double-headed black arrow between the lower-level representation of the critical region in the display and the object portion of the semantic network). A right/left side response is generated when the activity in the object NP exceeds threshold for one side of the display.

The green arrow labeled “R” emerging from the object NP in the semantic network indicates the participant’s right or left response; it’s shown in black to indicate that although the drift rate and threshold are affected by activity throughout the semantic network which affects activity within the NP representing the object, the R/L response must be dominated by recurrent activity between that NP and a lower-level representation of the object in the display.

Panel B depicts recurrent activity in the semantic network following presentation of valid labels (e.g., the label “lamp” for the object in the test display. The label pre-activates the semantic network associated with the object it denotes including the Object NP (dashed blue double headed arrows).

Double-headed black arrows indicate activation initiated by the object in the test display, as in A. Converging input to the Object NP from the semantic network and the input reduces the number of reentrant cycles necessary to generate the R/L response required by the task, thereby accounting for higher drift rates than control following valid labels.

In Panel C, dashed red lines illustrate activity in the semantic network following an invalid label denoting an object in a **different** superordinate-level category from the object in the test display (e.g., the label “bush” shown before the test display depicting a lamp). Semantic representations of bush throughout the semantic network are shown in red circles. Activation spreads throughout the semantic network, including to the Object NP for bush. The different colors of and spacing between the circles for lamp and bush illustrate that there is little overlap between the representations of objects in different superordinate-level categories. Accordingly, pre-activation of the semantic network by a label denoting an unrelated object in a different superordinate level category doesn’t affect the rate of evidence accumulation for the object in the test display. The greater activation of the semantic network because of this pre-activation may account for increased thresholds obtained in study 1 when valid and invalid labels are intermixed.

In Panel D, dashed purple lines illustrate activity in the semantic network following an invalid label denoting an object in the **same** superordinate-level category as the object in the test display (e.g., the label “drum” shown before the test display depicting a lamp). Semantic representations of drum throughout the semantic network are shown in purple circles. Activation spreads throughout the semantic network, including to the Object NP for drum. The similar colors of and spacing between the circles for drum and lamp illustrate overlap in the semantics networks of objects in the same superordinate-level category which produces noise. It is this noise that reduces drift rate and raises thresholds when same- superordinate level category invalid labels are used in study 2.

### Conflict or Noise?

Skocypec and Peterson (2022) proposed that in study 2 a conflict existed between the semantic networks activated by the invalid label and by the object in the test display. Our DDM model suggests a way to think about this conflict: There is more noise in the system in study 2 where invalid labels denote another object in the same superordinate-level category as the object in the test display. Greater noise lowers the signal-to-noise ratio (i.e., decreases drift rate). Indeed, we found that drift rate is lower following valid labels in study 2 than in study 1 and not higher for upright than inverted displays following invalid labels in study 2. Greater noise also increases threshold (i.e., more evidence must be accumulated from the display to differentiate the semantic networks activated by the label and the object). Indeed, we found threshold is higher following invalid than valid labels in study 2; no such difference was found in study 1. On this account, no additional conflict resolution process is necessary. Future research can adopt more complicated models to investigate if, when, and how a separate conflict resolution process occurs.

### Object detection entails semantic activation

The DDM results indicating that evidence thresholds are higher in labels-present studies than in labels-absent studies even following valid labels strongly implies that semantic activation plays a role in detection when labels are absent, supporting Skocypec and Peterson’s (2022) hypothesis that the detection of meaningful objects entails semantic activation rather than merely being affected by semantic activation. The finding of higher thresholds following same versus different superordinate-level invalid labels supports this hypothesis. Object detection does not occur until sufficient evidence has been accumulated to overcome noise in the semantic system.

### Sensitivity of the DDM

By analyzing RTs on both accurate and inaccurate trials, DDM modeling revealed some results that were not evident in Skocypec and Peterson’s (2022) separate analyses of participant’s mean accuracy and response times per condition, specifically that

1. evidence thresholds were raised when labels were present rather than absent, as discussed in the preceding section;
2. drift rate following valid labels was affected by the context of the type of invalid labels that were present on half of the trials; this context effect is consistent with prevalent sequential trial effects (e.g.,Fischer & Whitney, 2014; Pascucci et al., 2023). It would be interesting to explore the span of trials over which the effects of invalid labels persist.
3. drift rate is higher in the control condition for upright than inverted displays. Skocypec and Peterson were perplexed by observing that their control trial showed an orientation effect in accuracy but not in response time because it seemed that both effects were predicted by an explanation in terms of neural populations. Using RT obtained regardless of accuracy to assess the time needed to accumulate sufficient evidence for a response (drift rate) removes any questions and shows that drift rate is an excellent index of processing.

## Limitations of this Research

One limitation of our research is the model’s inability to explicitly distinguish whether the decision-making process involves one or multiple cognitive processes or stages. Traditional sequential sampling models like DDMs often assume a single integrated process of evidence accumulation (Bogacz et al., 2006; Ratcliff et al., 2016). However, this may oversimplify the neural processes involved, especially when considering recurrent neural processing loops in perception. In the current experiment, the perception of an object and its spatial location (R/L of a border) are not considered strictly linear or sequential processes but are presumed to involve iterative feedback loops. More specifically, detecting the object’s identity might be influenced by how its position is interpreted, and vice versa, with the brain constantly refining its interpretation as more information was processed and integrated in neurons in the LIP reflecting decision (e.g., Gold & Shadlen, 2007b; Ibos & Freedman, 2014). To resolve this limitation, future studies could incorporate more complex models that allow for multiple stages of decision-making, such as the hierarchical drift diffusion model or models with interactive competitive dynamics. These approaches can accommodate the parallel processing and feedback mechanisms inherent in neural computation.

Bornstein et al (2023) used a multi-stage DDM (MSDDM) to model their results. They manipulated the validity of pre-cues (50%, 60%, 70%, 80%) and the quality of the target stimuli, which were faces or scenes. Their results were best explained by a MSDDM, but their procedure was quite different from ours. The closest condition was a 50% valid cue condition followed by a high-resolution test display.

Unfortunately, they did not analyze their results from this condition, so their article is not informative regarding what might be learned from a MSDDM model of our results.

## Appendix

### Results for starting point

Study 1 valid label trials: orientation, *F*(1, 216) = 0.37, *p* = 0.544; study, *F*(1, 216) = 0.002, *p* = 0.962; interaction, *F*(1, 216) = 0.60, *p* = 0.439; invalid label trials: orientation, *F*(1, 216) = 2.31, *p* = 0.130; study, *F*(1, 216) = 0.30, *p* = 0.584; interaction, *F*(1, 216) = 0.00, *p* = 0.895. Study 2 valid label trials: orientation, *F*(1, 214) = 0.00, *p* = 0.996; study, *F*(1, 214) = 1.73, *p* = 0.190; and interaction, *F*(1, 214) = 2.05, *p* = 0.154; invalid label trials: orientation, *F*(1, 214) = 0.27, *p* = 0.603; study, *F*(1, 214) = 1.10, *p* = 0.297; interaction, *F*(1, 214) = 1.09, *p* = 0.299.

### Parameter recovery

In this analysis, we simulate behavioral data on our experiment with known parameter values and then fit this simulated data to see if the fit parameter values are close to the simulated parameter values. To generate parameters for the simulation, we used the same parameters that we obtained by fitting the full data set. In total, our simulated data set was approximately the same size as our real data set with 108 simulated participants each performing 72 trials in the control and experimental conditions; for the control experiment, we simulated data for 108 participants. For the experimental groups, we simulated data for only 108 participants rather than twice that number since the same model was used for study 1 and study 2. Thus, the simulated data approximated the number of participants tested in the experiment. The fitting procedure for parameter recovery was identical to that used for the real subjects.

In Figure 1 & 2, we plot the relationship between the recovered and simulated parameters for both the control and experimental models. To be noted, the parameters that were derived from the fitting process to the actual data were used to establish the boundaries for the parameters in the simulations.

**Figure 1.**
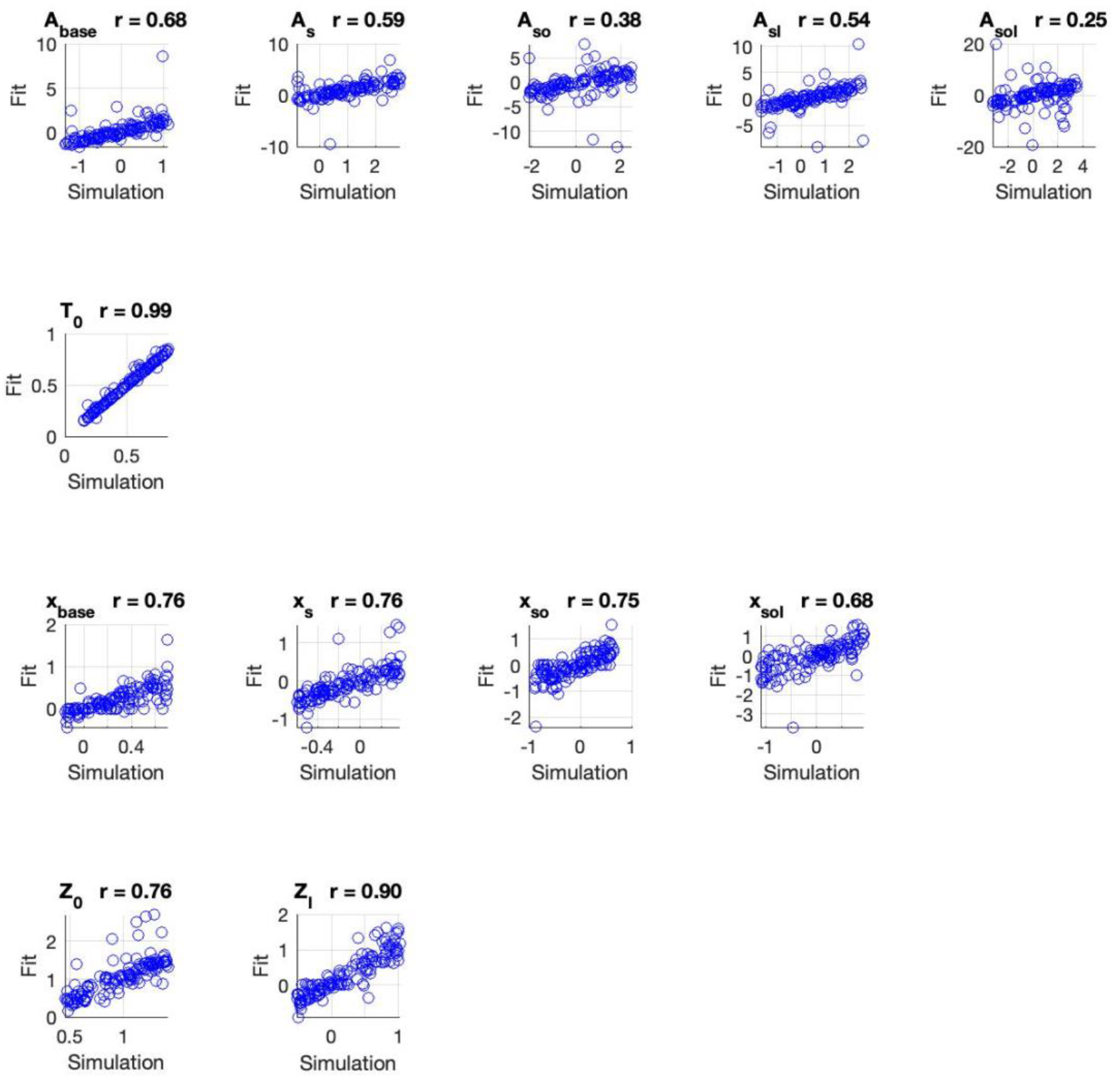
Parameter recovery for the full parameter model of the Experimental conditions with MLE fits. The value on the x axis represents simulation value, and value on y axis represents the value for the fitting process. Each row in the figure represents the components of up to 4 different parameters. Values of r on top of each axis represent the correlation coefficient. Note the scale of the y axis changes across graphs.

**Figure 2.**
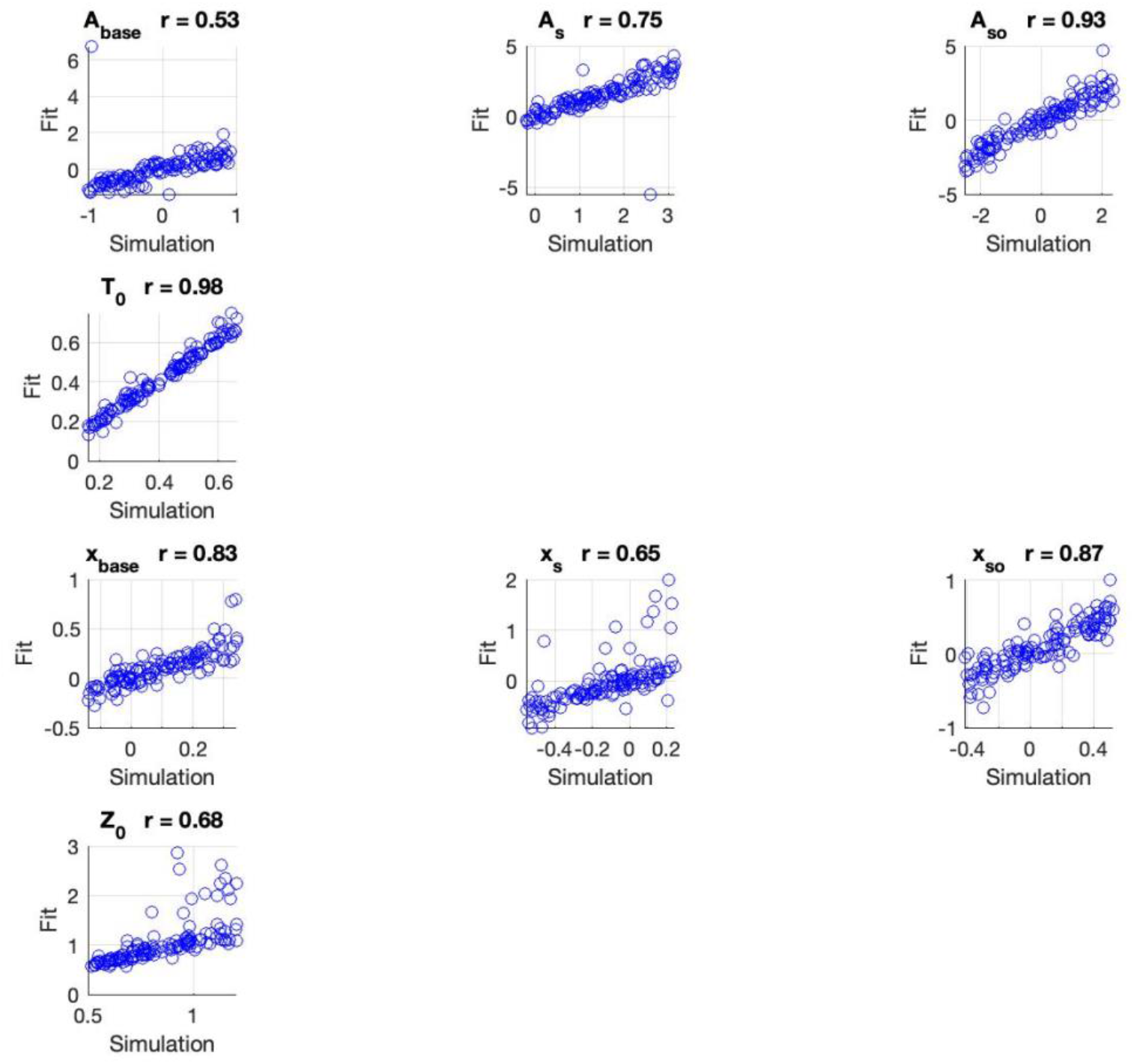
Parameter recovery for the model of the control conditions with MLE fits. Parameter recovery with MLE fits for the full parameter model. Axis and values of r have the same mean as in Figure 5, the only difference is that we have 8 instead of 12 parameterss for the control group.

Parameters that were recovered from this recovery that fell outside these predetermined boundaries were excluded. Despite this exclusion, the graphs demonstrate that the parameters were still very accurately recovered.

## References

1. Abdel Rahman, R., & Sommer, W. (2008). Seeing what we know and understand: How knowledge shapes perception. Psychonomic Bulletin and Review, 15(6). 10.3758/PBR.15.6.1055

2. Ashbridge, E., Perrett, D. I., Oram, M. W., & Jellema, T. (2000). Effect of image orientation and size on object recognition: Responses of single units in the macaque monkey temporal cortex. Cognitive Neuropsychology, 17(1–3). 10.1080/026432900380463

3. Bogacz, R., Brown, E., Moehlis, J., Holmes, P., & Cohen, J. D. (2006). The physics of optimal decision making: A formal analysis of models of performance in two-alternative forced-choice tasks. Psychological Review, 113(4). 10.1037/0033-295X.113.4.700

4. Bogacz, R., Hu, P. T., Holmes, P. J., & Cohen, J. D. (2010). Do humans produce the speed-accuracy trade-off that maximizes reward rate? Quarterly Journal of Experimental Psychology, 63(5), 863–891. 10.1080/17470210903091643

5. Bornstein, A.M., Aly, M., Feng, S.F. et al. Associative memory retrieval modulates upcoming perceptual decisions. Cogn Affect Behav Neurosci 23, 645–665 (2023). 10.3758/s13415-023-01092-6

6. Boutonnet, B., & Lupyan, G. (2015). Words jump-start vision: A label advantage in object recognition. Journal of Neuroscience, 35(25). 10.1523/JNEUROSCI.5111-14.2015

7. Brysbaert, M., & New, B. (2009). Moving beyond Kučera and Francis: A critical evaluation of current word frequency norms and the introduction of a new and improved word frequency measure for American English. Behavior research methods, 41(4), 977–990. 10.3758/BRM.41.4.977

8. Carr, T. H., McCauley, C., Sperber, R. D., & Parmelee, C. M. (1982). Words, pictures, and priming: On semantic activation, conscious identification, and the automaticity of information processing. Journal of Experimental Psychology: Human Perception and Performance, 8(6). 10.1037/0096-1523.8.6.757

9. Clarke, A., & Tyler, L. K. (2014). Object-specific semantic coding in human perirhinal cortex. Journal of Neuroscience, 34(14). 10.1523/JNEUROSCI.2828-13.2014

10. Costello, P., Jiang, Y., Baartman, B., McGlennen, K., & He, S. (2009). Semantic and subword priming during binocular suppression. Consciousness and Cognition, 18(2). 10.1016/j.concog.2009.02.003

11. Dils, A. T., & Boroditsky, L. (2010). Processing unrelated language can change what you see. Psychonomic Bulletin and Review, 17(6). 10.3758/PBR.17.6.882

12. Feng, S. F., Wang, S., Zarnescu, S., & Wilson, R. C. (2021). The dynamics of explore–exploit decisions reveal a signal-to-noise mechanism for random exploration. Scientific Reports, 11(1). 10.1038/s41598-021-82530-8

13. Fischer, J., Whitney, D. Serial dependence in visual perception. Nat Neurosci 17, 738–743 (2014). 10.1038/nn.3689

14. Flowers, C. S., Orsten-Hooge, K. D., Jannuzi, B. G. L., & Peterson, M. A. (2020). Normative data for an expanded set of stimuli for testing high-level influences on object perception: OMEFA-II. PLoS ONE, 15(8). 10.1371/journal.pone.0224471

15. Fodor, J. (1984). Observation Reconsidered. Philosophy of Science, 51(1). 10.1086/289162

16. Gayet, S., Van Der Stigchel, S., & Paffen, C. L. E. (2014). Breaking continuous flash suppression: Competing for consciousness on the pre-semantic battlefield. In Frontiers in Psychology (Vol. 5, Issue MAY). 10.3389/fpsyg.2014.00460

17. Gibson, B. S., & Peterson, M. A. (1994). Does Orientation-Independent Object Recognition Precede Orientation-Dependent Recognition? Evidence From a Cuing Paradigm. Journal of Experimental Psychology: Human Perception and Performance, 20(2). 10.1037/0096-1523.20.2.299

18. Gold, J. I., & Shadlen, M. N. (2007a). The neural basis of decision making. In Annual Review of Neuroscience (Vol. 30). 10.1146/annurev.neuro.29.051605.113038

19. Gold, J. I., & Shadlen, M. N. (2007b). The neural basis of decision making. In Annual Review of Neuroscience (Vol. 30<otherinfo>). 10.1146/annurev.neuro.29.051605.113038</otherinfo>

20. Ibos, G., & Freedman, D. J. (2014). Dynamic integration of task-relevant visual features in posterior parietal cortex. Neuron, 83(6). 10.1016/j.neuron.2014.08.020

21. Koffka. (1935). Principles of Gestalt Psychology (Chapter 1). In *Principles of Gestalt*.

22. Lositsky, O., Wilson, R. C., Shvartsman, M., & Cohen, J. D. (2015). A Drift Diffusion Model of Proactive and Reactive Control in a Context-Dependent Two-Alternative Forced Choice Task. *The Multi-Disciplinary Conference on Reinforcement Learning and Decision Making*.

23. Lupyan, G., & Clark, A. (2015). Words and the World: Predictive Coding and the Language- Perception-Cognition Interface. Current Directions in Psychological Science, 24(4). 10.1177/0963721415570732

24. Lupyan, G., & Ward, E. J. (2013). Language can boost otherwise unseen objects into visual awareness. Proceedings of the National Academy of Sciences of the United States of America, 110(35). 10.1073/pnas.1303312110

25. Maier, M., & Abdel Rahman, R. (2018). Native Language Promotes Access to Visual Consciousness. Psychological Science, 29(11). 10.1177/0956797618782181

26. Martin, C. B., Douglas, D., Newsome, R. N., Man, L. L. Y., & Barense, M. D. (2018). Integrative and distinctive coding of visual and conceptual object features in the ventral visual stream. ELife, 7. 10.7554/eLife.31873

27. Meteyard, L., Bahrami, B., & Vigliocco, G. (2007). Motion detection and motion verbs: Language affects low-level visual perception. Psychological Science, 18(11). 10.1111/j.1467-9280.2007.02016.x

28. Noorman, S., Neville, D. A., & Simanova, I. (2018). Words affect visual perception by activating object shape representations. Scientific Reports, 8(1). 10.1038/s41598-018-32483-2

29. Pascucci, D., Tanrikulu, Ö. D., Ozkirli, A., Houborg, C., Ceylan, G., Zerr, P., … & Kristjánsson, Á. (2023). Serial dependence in visual perception: A review. Journal of Vision, 23(1), 9–9. 10.1167/jov.23.1.9

30. Pedersen, M. L., Frank, M. J., & Biele, G. (2017). The drift diffusion model as the choice rule in reinforcement learning. Psychonomic Bulletin and Review, 24(4). 10.3758/s13423-016-1199-y

31. Perrett, D. I., Oram, M. W., & Ashbridge, E. (1998). Evidence accumulation in cell populations responsive to faces: An account of generalisation of recognition without mental transformations. Cognition, 67(1–2). 10.1016/S0010-0277(98)00015-8

32. Peterson, M. A. (2019). Past experience and meaning affect object detection: A hierarchical Bayesian approach. In Psychology of Learning and Motivation - Advances in Research and Theory (Vol. 70). 10.1016/bs.plm.2019.03.006

33. Peterson, M. A., & Gibson, B. S. (1994). Must figure-ground organization precede object recognition? An Assumption in Peril. Psychological Science, 5(5). 10.1111/j.1467-9280.1994.tb00622.x

34. Pinto, Y., van Gaal, S., de Lange, F. P., Lamme, V. A. F., & Seth, A. K. (2015). Expectations accelerate entry of visual stimuli into awareness. Journal of Vision, 15(8). 10.1167/15.8.13

35. Rabovsky, M., Sommer, W., & Abdel Rahman, R. (2012). Depth of conceptual knowledge modulates visual processes during word reading. Journal of Cognitive Neuroscience, 24(4). 10.1162/jocn_a_00117

36. Ratcliff, R. (1978). A theory of memory retrieval. Psychological Review, 85(2). 10.1037/0033-295X.85.2.59

37. Ratcliff, R., Smith, P. L., Brown, S. D., & McKoon, G. (2016). Diffusion Decision Model: Current Issues and History. In Trends in Cognitive Sciences (Vol. 20, Issue 4). 10.1016/j.tics.2016.01.007

38. Ratcliff, R., Thapar, A., & McKoon, G. (2004). A diffusion model analysis of the effects of aging on recognition memory. Journal of Memory and Language, 50(4). 10.1016/j.jml.2003.11.002

39. Rosenthal, V. (2006). Microgenesis, Immediate Experience and Visual Processes in Reading. In Seeing, Thinking and Knowing. 10.1007/1-4020-2081-3_11

40. Samaha, J., Boutonnet, B., Postle, B. R., & Lupyan, G. (2018). Effects of meaningfulness on perception: Alpha-band oscillations carry perceptual expectations and influence early visual responses. Scientific Reports, 8(1). 10.1038/s41598-018-25093-5

41. Sander, F. (2006). Structure, totality of experience, and Gestalt. In Psychologies of 1930. 10.1037/11017-010

42. Skocypec, R. M., & Peterson, M. A. (2022). Semantic Expectation Effects on Object Detection: Using Figure Assignment to Elucidate Mechanisms. Vision (Switzerland), 6(1). 10.3390/vision6010019

43. Stein, T., & Peelen, M. V. (2015). Content-specific expectations enhance stimulus detectability by increasing perceptual sensitivity. Journal of Experimental Psychology: General, 144(6). 10.1037/xge0000109

44. Stokes, M., Thompson, R., Nobre, A. C., & Duncan, J. (2009). Shape-specific preparatory activity mediates attention to targets in human visual cortex. Proceedings of the National Academy of Sciences of the United States of America, 106(46). 10.1073/pnas.0905306106

45. Tavares, G., Perona, P., & Rangel, A. (2017). The attentional Drift Diffusion Model of simple perceptual decision-making. Frontiers in Neuroscience, 11(AUG). 10.3389/fnins.2017.00468

46. Weller, P. D., Rabovsky, M., & Abdel Rahman, R. (2018). Semantic knowledge enhances conscious awareness of visual objects. Journal of Cognitive Neuroscience, 31(8). 10.1162/jocn_a_01404

47. Wilson, R. C., & Collins, A. G. E. (2019). Ten simple rules for the computational modeling of behavioral data. ELife, 8. 10.7554/eLife.49547

